# Expansion of tissue-resident CD8+ T cells and CD4+ Th17 cells in the nasal mucosa following mRNA COVID-19 vaccination

**DOI:** 10.1101/2021.05.07.442971

**Authors:** Aloysious Ssemaganda, Huong Mai Nguyen, Faisal Nuhu, Naima Jahan, Catherine M. Card, Sandra Kiazyk, Giulia Severini, Yoav Keynan, Ruey-Chyi Su, Hezhao Ji, Bernard Abrenica, Paul J. McLaren, T. Blake Ball, Jared Bullard, Paul Van Caeseele, Derek Stein, Lyle R. McKinnon

**Affiliations:** Department of Medical Microbiology and Infectious Diseases, University of Manitoba, Winnipeg, MB, Canada; JC Wilt Infectious Diseases Research Centre, National Microbiology Laboratory, Public Health Agency of Canada, Winnipeg, MB, Canada; Cadham Provincial Laboratory, Winnipeg, MB, Canada; Department of Pediatrics & Child Health, University of Manitoba, Winnipeg, MB, Canada; Centre for the AIDS Programme of Research in South Africa (CAPRISA), Durban, South Africa

**Author notes:** **Corresponding Author:** Dr. Lyle McKinnon, Department of Medical Microbiology and Infectious Diseases, 504-745 Bannatyne Ave, University of Manitoba, Winnipeg, MB, Canada. R3E 0J9.

## Abstract

Vaccines against SARS-CoV-2 have shown high efficacy in clinical trials, yet a full immunologic characterization of these vaccines, particularly within the upper respiratory tract, remains lacking. We enumerated and phenotyped T cells in nasal mucosa and blood before and after vaccination with the Pfizer-BioNTech COVID-19 vaccine (n =21). Tissue-resident memory (Trm) CD8+ T cells expressing CD69+CD103+ expanded ∼12 days following the first and second doses, by 0.31 and 0.43 log_10_ cells per swab respectively (p=0.058 and p=0.009 in adjusted linear mixed models). CD69+CD103+CD8+ T cells in the blood decreased post-vaccination. Similar increases in nasal CD8+CD69+CD103-T cells were observed, particularly following the second dose. CD4+ Th17 cells were also increased in abundance following both doses. Following stimulation with SARS-CoV-2 spike peptides, CD8+ T cells increased expression of CD107a and CD154. These data suggest that nasal T cells may be induced and contribute to the protective immunity afforded by this vaccine.

## Main text

The COVID-19 pandemic has led to substantial mortality and caused major global disruption. At least six vaccines have demonstrated efficacy in preventing severe disease associated with SARS-CoV-2 infection^1^. While this is an impressive achievement guided by early immunogenicity data, a full understanding of how these novel vaccines interact with the host immune system is lacking. This is particularly the case in tissues such as the upper respiratory tract (URT), which are often more challenging to sample than peripheral blood^2^. Here we focus on characterizing nasal mucosal T cells in healthy individuals pre- and post-vaccination with the BNT162b2 mRNA-nanoparticle vaccine developed by Pfizer-BioNTech^3^.

Natural history studies of SARS-CoV-2 infection demonstrate significant cell-mediated and humoral immune responses^4-7^, which in combination lead to a substantial decrease in re-infection risk^8^. These responses can be readily detected for >8 months post-infection^9^, and appear to have a long half-life. In non-human primates, CD8+ T cell depletion led to increased susceptibility to re-infection^10^. Studies that assess the quality and quantity of immune responses induced by vaccination in humans are beginning to emerge^11-16^, adding depth to what was generated in early phase clinical trials. While these studies suggest a high magnitude of humoral and cell-mediated immunity to SARS-CoV-2^17,18^, less is known regarding tissue and mucosal immunity induced by these vaccines, particularly in the URT, the primary site of viral entry.

We developed an *ex vivo* flow cytometry-based assay to enumerate and profile immune cells isolated from nasopharyngeal (NP) swabs used for SARS-CoV-2 diagnostic testing amongst healthy volunteers. We pilot-tested several types of NP swabs to gauge optimal immune cell recovery. Strikingly, CD45+ immune cells were recovered from a certain type of swab (Flexible minitip flocked swab from BD), while CD326+ epithelial cells were the main cell type recovered from two other swabs that we tested, including the Copan FLOQ swab (**Figure 1A &B**). In a pilot study (n=8), a median of 3,082 (IQR: 2,351-6,168) CD45+ cells were recovered per BD swab, ∼80% of which were CD3+ T cells. Using CD69 and CD103 as markers of tissue-resident memory (Trm) T cells^19,20^, virtually all CD8+ (>90%) and variable proportions of CD4+ T cells were Trm (10-80%; **Figure 1C**).

**Figure 1:**
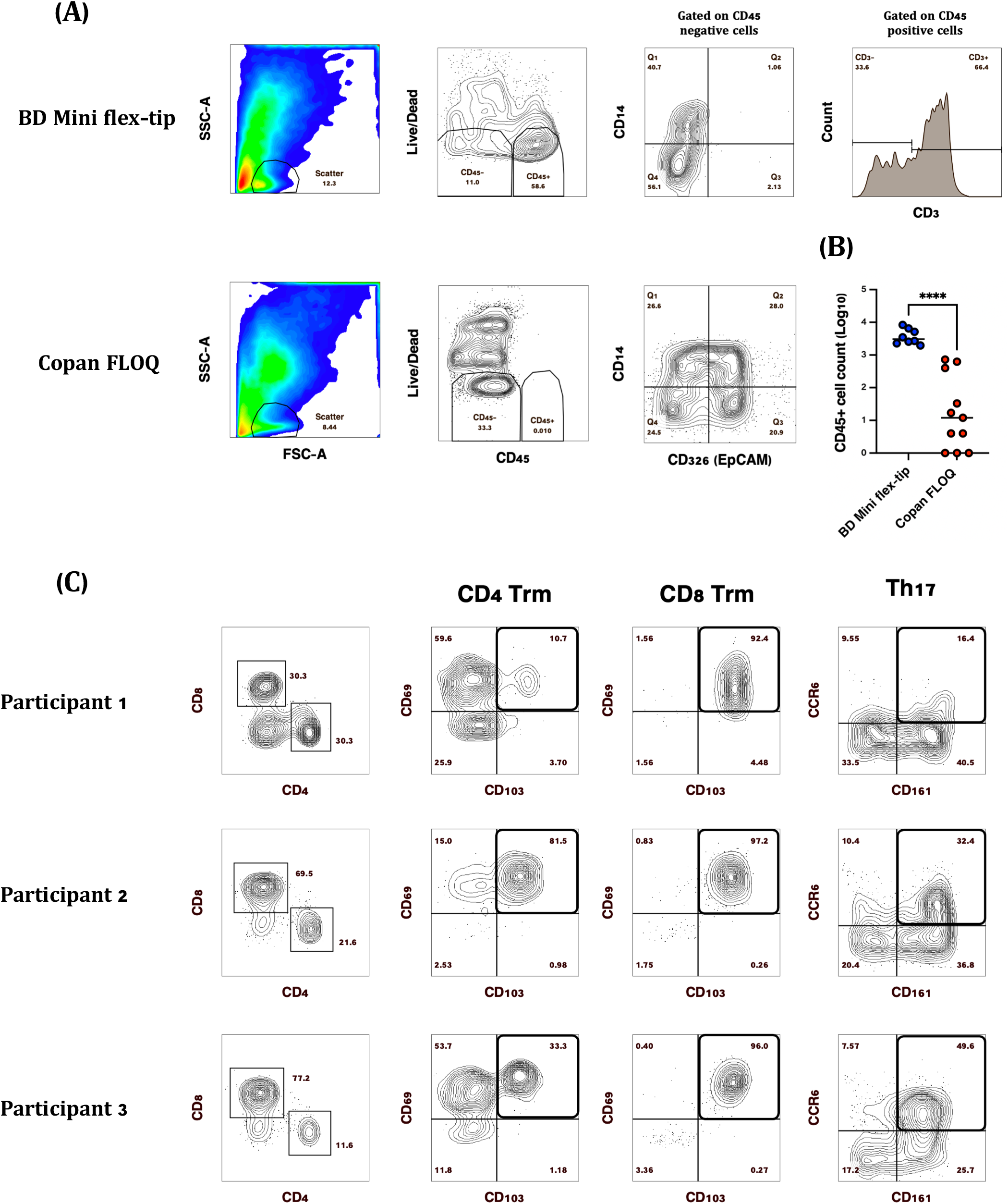
Comparison of SARS-CoV-2 diagnostic testing nasopharyngeal (NP) swabs for immune cell recovery. Representative plots of CD45+ nasal immune cells isolated from (**A**) BD Mini Flex-tip swabs, majority of which (∼40-90%) were CD3+ and Copan FLOQ swabs from which epithelial (CD326 (EpCAM+) cells were the main cell type recovered. (**B**) Swab brand comparison of CD45+ immune cell recovery. **(C)** Representative plots of CD3+ T cell subsets isolated from BD Mini-tip flexible swabs. Majority of CD4+ and CD8+ T cells were Trm based on the expression of CD69 and CD103, while 16-50% of CD4 T cells were Th17 based on the expression of CD161 and CCR6. (****P>0.0001)

We enrolled healthy volunteers scheduled to receive the BNT162b2 mRNA vaccine (n=21, **Supplemental Table 1**), and collected baseline, pre-vaccine samples followed by peak response samples ∼12 days after the first (visit 2) and second (visit 3) doses. Participants were a median age of 40 (IQR: 31-50) and predominantly female (14/21). Most participants (19/21) received influenza vaccination in the previous few months, and two participants had prior COVID-19 infection. The median body mass index (BMI) was 24 (IQR: 22-29). Reporting of underlying medical conditions included allergies and/or asthma (n=6), type II diabetes (n=2), hypertension (n=2), and autoimmunity (n=1).

We analyzed both frequency of parent populations and total cell abundance, the latter by maximizing acquisition of cells from each swab; measuring abundance is critical as cell abundance can vary within mucosa across orders of magnitude^21,22^. We observed significant increases in nasal CD8+ Trm at both peak vaccine response time points (**Figure 2A**). In a linear mixed model adjusted for age, sex, BMI, prior COVID-19, influenza vaccination, and self-reported vaccine side effects, the number of CD8+ Trm increased by 0.31 and 0.44 log_10_ cells per nasal swab following the first and second vaccine dose, respectively (p=0.058 and p=0.009). No increases in CD4+ Trm defined by CD103 and/or CD69 expression were observed **(Figure 2B)**. The absolute number of nasal CD8+CD69+CD103-T cells also increased at visit 2 and particularly at visit 3, by 0.06 and 0.48 log_10_ cells/swab, respectively (adjusted p=0.7 and p=0.004 for visits 2 and 3). The proportion of nasal CD8+ T cells that were Trm also increased, although not significantly (p=0.09 from visit 2 versus baseline, **Supplemental Figure 1A-B**). Interestingly, the low proportion of CD69+CD103+ T cells in the blood significantly declined at the peak vaccine response time points, suggesting that these cells may be migrating out of the blood into tissues **(Figure 2C-D)**. We also observed that CD4+ T cells co-expressing the Th17 markers CD161 and CCR6 increased by 0.40 and 0.45 log_10_ cells/nasal swab at the peak vaccine response time points compared to baseline (adjusted p=0.017 and p=0.008, respectively) **(Figure 2E)**. No increases in nasal Tfh cells were observed following either vaccine dose **(Figure 2F)**. Similar increases were observed in analyses stratified by age and sex (**Supplemental Table 2**), and excluding participants with prior COVID-19 infection did not meaningfully change any outcomes.

**Figure 2:**
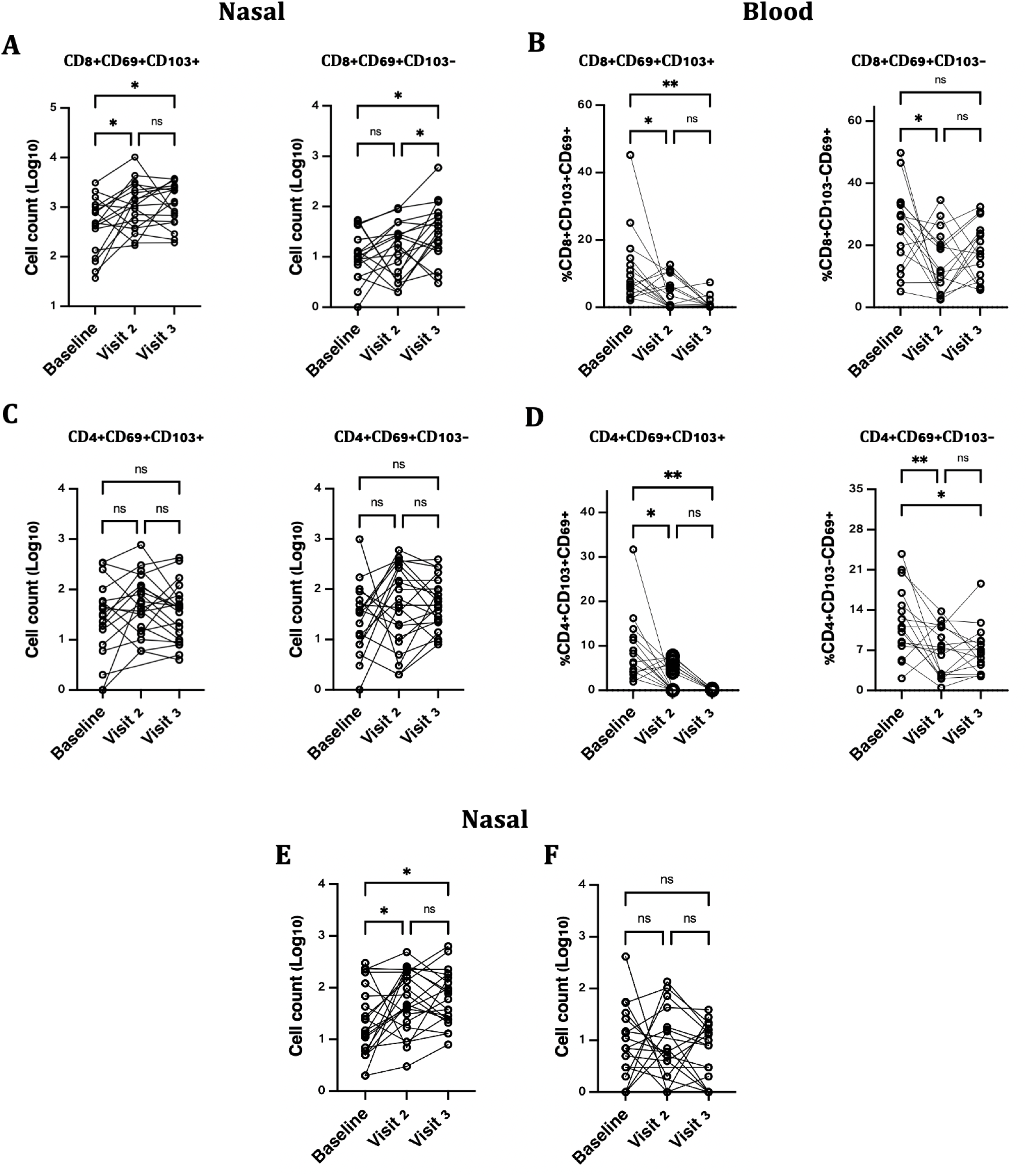
Nasal cell counts per NP swab and peripheral blood frequencies of T cell subsets post SARS-CoV-2 vaccination. (A) Nasal CD8+CD69+CD103+ Trm significantly increased following the first and second vaccine dose while CD8+CD69+CD103-T cells increased significantly following the second vaccine dose. (B) No changes in nasal CD4 +CD69+CD103+ Trm (C) In blood, CD8+CD69+CD103+ Trm and (D) CD4+CD69+CD103+ decreased significantly at visit 2 and 3 compared to baseline. (E) Significant increases in Th17 (CD4+CD161+CCR6+) T cells at peak vaccine response time points while (E) no changes were observed in Tfh (CD4+CXCR5+PD1+) T cells.(* P>0.01, **P>0.001, ns: non-significant).

Studies of other vaccines suggest that peak vaccine-elicited T cells can be captured by activation markers such as Ki67, HLA-DR, and CD38, which peak at 10-14 days following yellow fever and smallpox vaccination^23^. In our study, we did not observe any changes in the proportion of CD4+ or CD8+ T cells co-expressing HLA-DR and CD38, or any upregulation of Ki-67, in either nasal or blood samples (**Supplemental Figure 2**).

We further analysed our flow data using the t-distributed stochastic neighbour embedding (tSNE), a dimensionality reduction method. Several substantial shifts in CD45+ nasal immune cell clusters were evident from baseline to visit 2, and from visit 2 to 3 **(Supplemental Figure 3A)**. These included a step-wise shift in CD8+ T cell populations across the three time points, but also shifts in CD3-and CD3+CD4-CD8-cells. In support of this, the abundance of CD45+, CD3+, and CD8+ T cells all increased at visit 3, compared to baseline **(Supplementary Figure 3B-G)**. While it was difficult to distinguish which phenotypic markers flow panel explained the observed divergent clustering patterns, these data suggest that multiple subsets of nasal immune cells may increase in number following COVID-19 vaccination.

**Figure 3:**
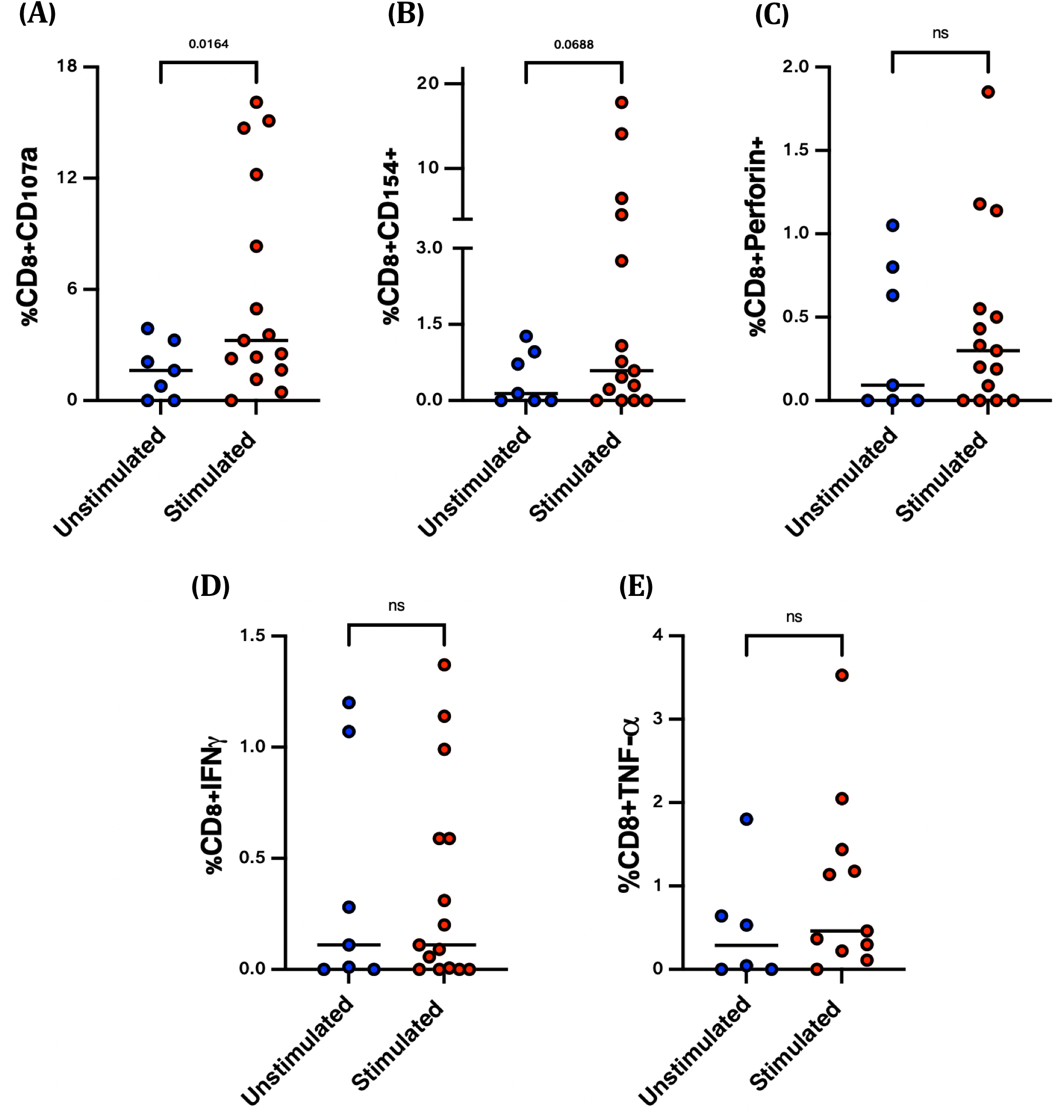
SARS-CoV-2 spike-specific CD8+ T cell responses ∼2 months post-vaccination. Nasal immune cells isolated from individuals, approximately 2 months post second vaccine dose, were stimulated (n=15) with overlapping spike peptides (Red dots) or left unstimulated (n=7) (Blue dots). Frequencies of **(A)** CD107a, **(B)** CD154, **(C)** Perforin, **(D)** TNF-α and **(E)** IFN-γ antigen-specific CD8+ T cell responses. (ns: non-significant).

To further assess the antigen-specificity of nasal T cells, we enrolled vaccinated individuals at 2 months following their second Pfizer-BNT dose, and either stimulated the NP swab cells with overlapping SARS-CoV-2 spike protein peptide pools (n=15), or left cells in media unstimulated (n=7). Stimulated samples had increased expression of the cytotoxic marker, CD107a (p=0.016) and the activation marker CD154 (p=0.069, **Figure 3**). Similar trends were observed for TNF-α and perforin, though these were not statistically significant.

To understand the relationship between antibody titres and nasal T cells, we quantified levels of SARS-CoV-2 spike-specific IgG antibodies in plasma. Antibody titres increased at visit 2 (median titre 163, IQR: 67-376), and more dramatically at visit 3 (median 2,185, IQR: 826-3,652; **Figure 4A**). Only one of two participants with prior COVID-19 was positive for SARS-CoV-2 nucleocapsid IgG (**Figure 4B**). We next correlated antibody titres to the number of nasal CD8+ Trm and CD4+ Th17 at visit 3. Nasal CD8+ Trm and CD4+ Th17 increases were similar in participants with varying increases in spike titres, with no correlations observed (**Figure 4C-D**). These data suggest that tissue T cell responses to vaccination may be independent of the antibody response and may provide an additional layer of immunity against SARS-CoV-2 infection.

**Figure 4:**
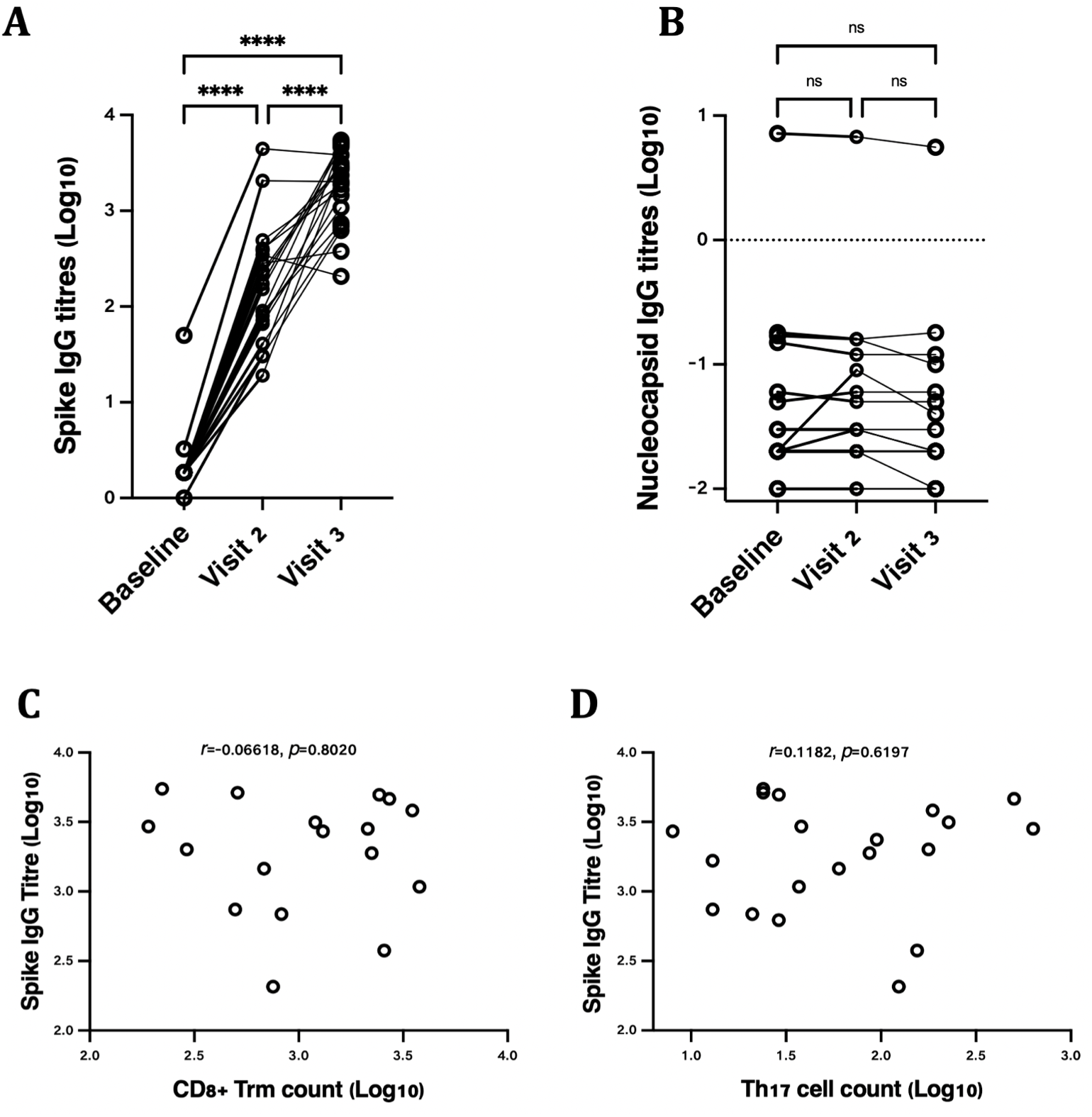
Plasma IgG antibody titres following SARS-COV-2 vaccination (A) spike-specific IgG titres and (B) nucleocapsid specific IgG titres. (C) Correlation of nasal CD8+ Trm and CD4+ Th17 T cell counts with spike antibody titres at visit 3, ∼12 days following the second SARS-COV-2 vaccine dose (****p<0.0001, ns: non-significant).

Trm are a vital form of surveillance in mucosal tissues, where they can mount rapid immune responses to pathogens upon re-exposure. In animal models, T cells at peak activation up-regulate homing markers that distribute these cells to relevant tissues^24^, where a high proportion is retained for extended periods as a form of sentinel surveillance against re-infection^25^. Trm have been more difficult to study in humans, given challenges in sampling the hard-to-reach tissues and in proving residency^26,27^. Our data suggest that certain types of NP swabs commonly used for COVID-19 diagnostics are effective in recovering immune cells from nasal tissue. Multiple populations of nasal CD4+ and CD8+ T cells increased in frequency by ∼0.5 orders of magnitude following vaccination with the BNT162b2 mRNA vaccine. This is a critical observation, as most human mucosal COVID-19 studies have focused either on antibodies^28^ or lung T cells^29,30^; while a tissue population, the latter may not represent the front-line defense against infection. It has been hypothesized that Trm may be the most effective form of T cell response against SARS-CoV-2 infection^31^, capable of preventing viral dissemination beyond the URT, where more virus-induced damage can occur^32^. Data from mice have suggested that parenteral vaccination induces nasal Th17 Trm that protect against bacterial colonization^33,34^. However, our study is the first, to our knowledge, to suggest nasal T cells may be induced by parenteral vaccination against COVID-19.

Our study has some limitations. We enrolled a relatively small sample size, restricting our ability to extensively stratify our data. The age range does not include the elderly, a group most in need of protection by COVID-19 vaccination^35^. The immune cell recovery from NP swabs, while sufficient for phenotyping and enumeration, is challenging for extensive functional characterization, in particular in assays that required splitting samples for paired analyses.

In summary, nasal T cell studies are feasible in humans, but this is highly dependent on the type of NP swab that is used. A high proportion of nasal T cells express markers of tissue residency, and these cells increase significantly in number and function following SARS-CoV-2 mRNA immunization. Further work should be performed to understand the role that local T cells may play in the protection against SARS-CoV-2 infection.

## Acknowledgements

This work was funded by the Bill and Melinda Gates Foundation and the Canadian Institutes of Health Research. Special thanks to the study volunteers whose time and willingness to provide repeated specimens made this project possible.

## Author contributions

AS, DS, LRM: Conceptualized the study, designed the experiments and analysed data. LRM, TBB, JB, PVC: Overall study supervision and coordination.

CMC, SK, YK, RS, HJ, PJM, TBB, JB, PVC, DS: Provided intellectual input into study design. NJ, GS, BA: Enrolled study participants, carried out sampling and/or collected and entered data.

AS, HMN, FN, DS: Performed experiments.

AS, LRM: Wrote the initial draft manuscript.

All authors reviewed and approved the submitted manuscript.

---

**Supplemental Table 1.** Participant demographics and health status.

| Variable | Study participants (n= 21) |
| --- | --- |
| Age (median, IQR) | 40 (31-50) |
| Sex, female | 67% (14/21) |
| Body mass index (BMI, median, IQR) | 24.4 (21.5-28.9) |
| Received 2020 fall flu vaccine | 90% (19/21) |
| Diagnosed with COVID-19 in 2020 | 10% (2/21) |
| Self-reported underlying medical conditions | Asthma/allergy (6/21)<br>Type 2 diabetes (2/21)<br>Hypertension (2/21)<br>Autoimmunity (1/21) |
| Dose 2 side effects | None (5/21)<br>Sore arm (3/21)<br>Moderate effects including nausea, fever, influenza-like symptoms (13/21) |

**Supplemental Table 2.** Age and Sex-stratified analysis of CD8+ Trm and Th17 cells following SARS–CoV-2 vaccination

| Strata |  | Outcome* (V3 vs baseline; beta & 95% CI) |  |  |
| --- | --- | --- | --- | --- |
|  |  | CD8+<br>CD69+CD103+ | CD8+<br>CD69+CD103- | CD4+<br>CCR6+CD161+ |
| **Entire cohort** |  | 0.43 (0.11-0.64) | 0.48 (0.16-0.81) | 0.45 (0.12-0.78) |
| **Sex** | Males (n=7) | 0.57 (0.08-1.07) | 0.39 (-0.11-0.90) | 0.38 (-0.07-0.83) |
|  | Females (n=14) | 0.39 (0.00-0.78) | 0.51 (0.11-0.90) | 0.49 (0.08-0.90) |
| **Age** | ≤35 years of age (n=8) | 0.21 (-0.30-0.71) | 0.22 (-0.18-0.62) | 0.42 (-0.14-0.98) |
|  | >35 to <50 years of age (n=6) | 0.85 (0.68-1.02) | 1.00 (0.56-1.44) | 0.55 (0.18-0.91) |
|  | ≥50 years of age (n=7) | 0.45 (-0.06-0.95) | 0.42 (-0.09-0.94) | 0.41 (-0.13-0.95) |

**Supplementary Table 3.**
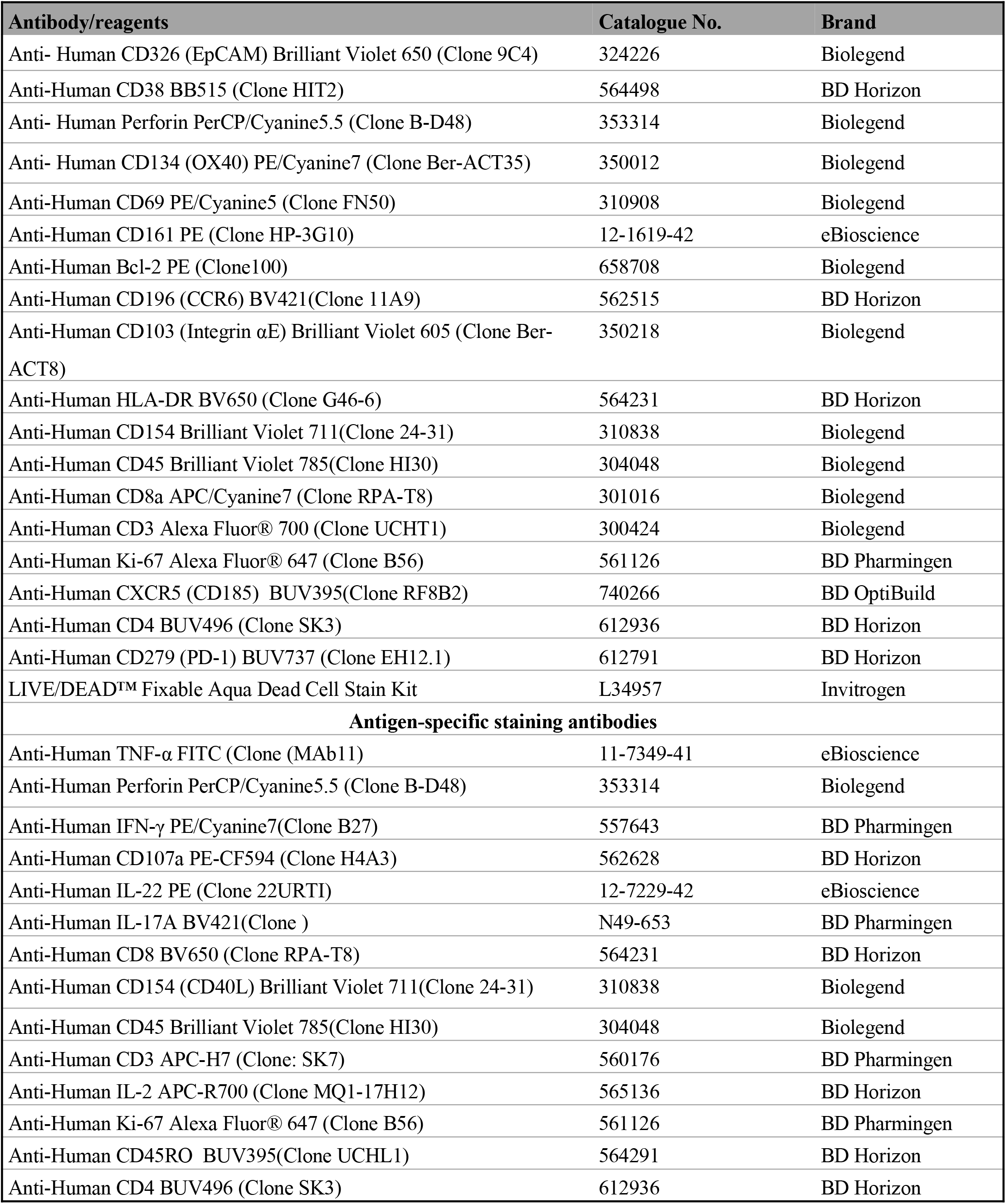
List of flow cytometry antibodies/reagents

**Supplementary Figure 1:**
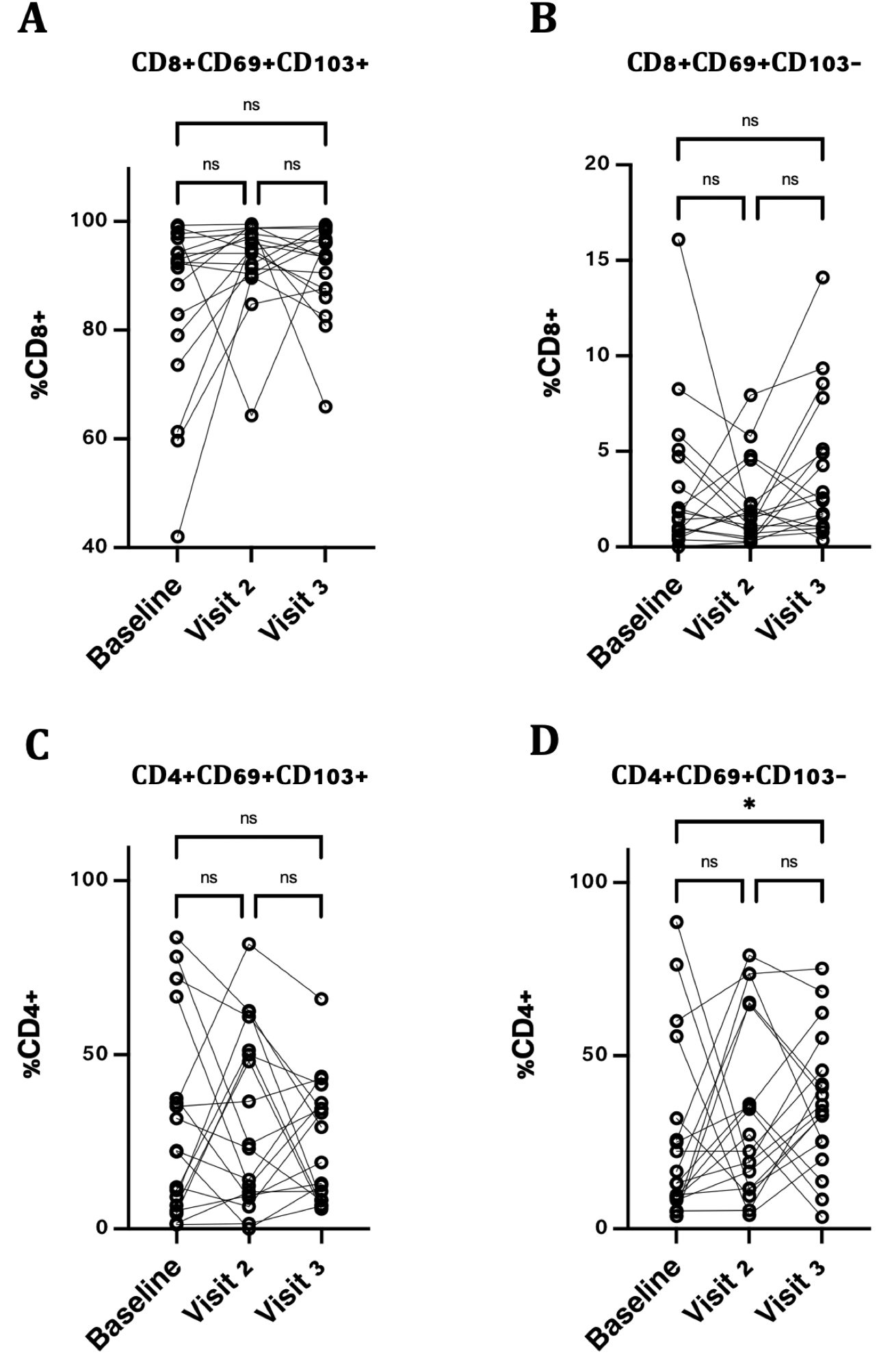
Frequencies of nasal CD4 and CD8 Trm T cells post SARS-COV-2 vaccination. Proportions of (A) CD8+CD69+CD103+, (B) CD8+CD69+CD103-, (C) CD4+CD69+CD103+Trm and (D) CD4+CD69+CD103+ Trm T cells (* P>0.01, ns: non-significant).

**Supplementary Figure 2.**
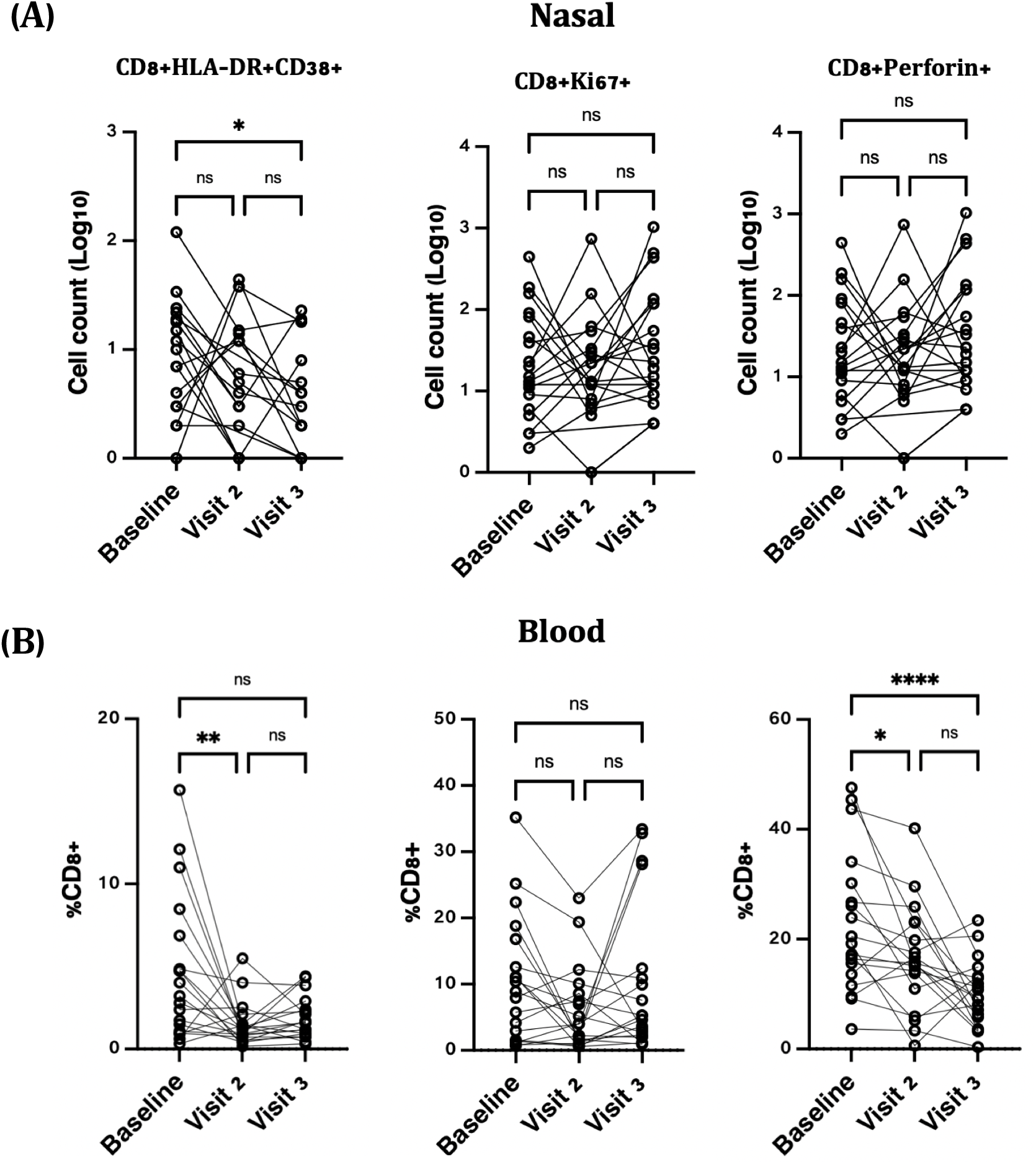
CD8+ T cell activation defined by expression of HLA-DR and CD38, proliferation, based on the expression of Ki-67 and cytotoxicity, defined by the expression of perforin in cells isolated from (A) nasopharyngeal swabs and (B) peripheral blood amongst vaccinated individuals (n=21) following SARS-COV-2 vaccination (* p<0.01, **p<0.001, ****p<0.0001, ns: non-significant).

**Supplementary Figure 3.**
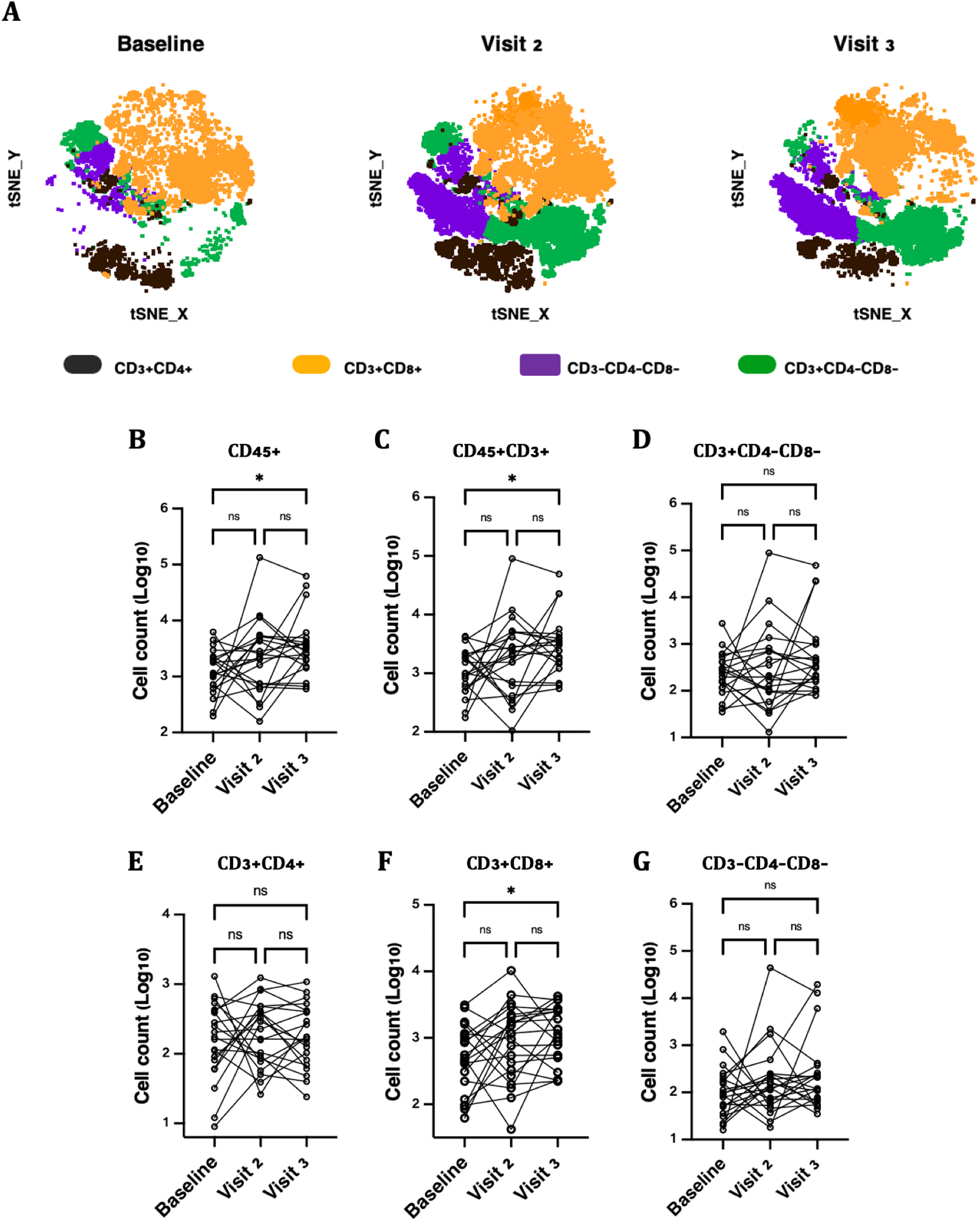
Nasal immune cell clustering and cell counts per swab following SARS-COV-2 vaccination. (A) t-distributed stochastic neighbour embedding (tSNE) plots of integrated nasal CD45+ immune cells showing shifts in CD3+CD4+, CD3+CD8+, CD3-CD4-CD8- and CD3+CD4-CD8-immune cell clusters from vaccinated individuals at baseline, visit 2 and visit 3. (B) CD45+ (C) CD45+CD3+ and (F) CD3+CD8+ cells were significantly increased following the 2nd vaccination dose while no changes were observed for (D) CD3+CD4-C8-, (E) CD3+CD4+ and (G) CD3-CD4-CD8-immune cell subsets (* P>0.01, ns: non-significant).

## Supplemental Methods

### Study participants and ethics board approval

Study volunteers (n=29) included staff enrolled at the Cadham Provincial Laboratory in Winnipeg, with ethics approval from the University of Manitoba Health Research Ethics Board (HREB) and the Public Health Agency of Canada (PHAC). Informed consent was obtained and a comprehensive socio-demographic and health assessment questionnaire administered to all study participants.

### Sample collection and processing

In pilot work, we compared the BD Flex-mini and Copan FLOQ swabs. In the vaccine study participants, matched diagnostic pharyngeal (NP) swabs (BD Flex-mini) and peripheral blood samples (BD EDTA Vacutainer) were collected prior to vaccination (baseline), ∼12 days after the first vaccine dose (visit 2), and ∼12 days after the second dose (visit 3). Nasal cells were isolated from NP swabs which were inserted into the nose, rotated once (360°) and placed into a tube containing 3ml of viral transport medium (VTM) and placed on ice for processing within 2 hours of collection. Once in the laboratory, swabs were vigorously vortexed to dissociate the cells and mucus and centrifuged for 10 min at 1600rpm. NP swabs were then rinsed with PBS 2% FBS, and the cell suspension filtered through 100µm nylon cell strainer (Becton Dickinson) fitted into a 50 ml tube. Cells were washed twice prior to flow cytometry staining. Peripheral blood mononuclear cells (PBMC) were isolated by ficoll density gradient centrifugation using SepMate™ 50 (STEMCELL ™ technologies) following the manufacturer’s instructions.

### Flow cytometry

Cells were stained with viability exclusion dye, together with a panel of pre-titrated surface antibodies (**Supplementary Table 1**) in PBS 2% FBS on ice for 30 min. The NP swab optimization experiment (**Figure 1**) included anti-human CD326 (EpCAM) (Biolegend, Clone 9C4) in the surface staining cocktail to stain for epithelial cells. Cells were subsequently washed, permeabilized in fixation/permeabilization solution (eBiosciences) and stained for Ki*-*67 and perforin. The cells were washed again and resuspended in perm/wash buffer (eBioscience). Data were acquired using a BDLSR Fortessa flow cytometer (BD Biosciences) and analyzed using FlowJo™ version 10.7.1 (Becton Dickinson Life Sciences).

### Dimensionality reduction, visualization and clustering

FlowJo™ version 10.7.1 (Becton Dickinson Life Sciences), was utilized to dimensionally reduce and to interrogate immune cell clusters and phenotypes in cells isolated from nasal swabs. Briefly, manual gating was performed to exclude debris, doublets and dead cells. Live CD45+ cells from all participants (n=21) at baseline, visit 2, and visit 3, were integrated into a single file which was dimensionally reduced and visualized using t-distributed stochastic neighbour embedding (tSNE), following the default parameter settings. FlowSOM algorithm (Version 2.6) was used to create cluster populations from the dimensionally reduced data based on their similarity and subsequently segregated based on relative abundance and phenotype using ClusterExplorer (Version 1.5.9).

### Antigen-specific stimulation and flow cytometric staining of nasal cells

Approximately 2 months post-2^nd^ vaccination, participants (n=15) were enrolled. Freshly isolated nasal cells from (n=11) were stimulated for six hours at 37 °C / 5% CO_2_ with SARS–CoV-2 spike peptide pools S, S1 and S+ containing 15-mer sequences overlapping by11 amino acids (PepTivator^®^, Miltenyi Biotec), encompassing the complete spike protein sequence (GenBank MN908947.3) at a final concentration of 1µg/mL per peptide, in the presence of Golgi-Plug (BD Biosciences), Golgi-Stop (BD Biosciences) and anti–CD107a (clone H4A3, BD Biosciences). As a control for this experiment, n=4 participants’ cells were left unstimulated. Following stimulation, cells were washed, stained with Live/Dead ™ Fixable Aqua viability dye (Invitrogen) for 30 mins on ice. Cells were subsequently washed, permeabilized with fixation/permeabilization solution (eBiosciences) and stained in a cocktail of pre-titrated monoclonal antibodies (**Supplementary Table 1**). Cells were then washed, acquired using a BDLSR Fortessa (BD Biosciences) and analyzed using FlowJo™ version 10.7.1 (Becton Dickinson Life Sciences).

### Plasma antibody testing

Concentrations of SARS-CoV-2-specific IgG were determined using commercial assays. At visit 3, plasma was diluted 1:10 because of the high magnitude of the responses. IgG recognizing the SARS-CoV-2 spike protein was quantified using the Diasorin Trimeric Spike ELISA, while nucleocapsid antibodies were measured using the ELISA from Abbott.

### Statistics

Clinical and demographic data are presented as proportions or medians and interquartile ranges. Linear mixed models were used to compare changes in log10 transformed immune cell abundance (in nasal mucosa, typically cells/swab; in blood, % of CD3+CD4+ or CD8+ T cells). In these models, visit as a categorical variable was the main predictor, with visit 2 and 3 compared to baseline. Multivariate models were adjusted for a range of covariates. Raw p-values, beta coefficient and 95% confidence intervals were reported. Statistical analyses were performed using SPSS v. 27.0 while all the graphing and was performed using Prism version 9 (GraphPad Software, LLC).

